# MRCZ – A proposed fast compressed MRC file format and direct detector normalization strategies

**DOI:** 10.1101/116533

**Authors:** Robert A. McLeod, Ricardo Diogo Righetto, Andy Stewart, Henning Stahlberg

## Abstract

The introduction of high-speed CMOS detectors is fast marching the field of transmission electron microscopy into an intersection with the computer science field of big data. Automated data pipelines to control the instrument and the initial processing steps are imposing more and more onerous requirements on data transfer and archiving. We present a proposal for expansion of the venerable MRC file format to combine integer decimation and lossless compression to reduce storage requirements and improve file read/write times by >1000 % compared to uncompressed floating-point data. The integer decimation of data necessitates application of the gain normalization and outlier pixel removal at the data destination, rather than the source. With direct electron detectors, the normalization step is typically provided by the vendor and is not open-source. We provide robustly tested normalization algorithms that perform at-least as well as vendor software. We show that the generation of hot pixels is a highly dynamic process in direct electron detectors, and that outlier pixels must be detected on a stack-by-stack basis. In comparison, the low-frequency bias features of the detectors induced by the electronics on-top of the active layer, are extremely stable with time. Therefore we introduce a stochastic-based approach to identify outlier pixels and smoothly filter them, such that the degree of correlated noise in micrograph stacks is reduced. Both *a priori* and *a posteriori* gain normalization approaches that are compatible with pipeline image processing are discussed. The *a priori* approach adds a gamma-correction to the gain reference, and the *a posteriori* approach normalized by a moving average of time-adjacent stacks, with the current stack being knocked-out, known as the KOMA (knock-out moving average) filter. The combination of outlier filter and KOMA normalization over ~25 frames can reduce the correlated noise in movies to nearly zero. Sample libraries and a command-line utility are hosted at *github.com/em-MRCZ* and released under the BSD license.

## 1 Introduction

The introduction of CMOS-based direct electron detectors for transmission electron microscopy greatly improved the duty cycle to nearly 100 % compared to traditional slow-scan CCD detectors. The high duty-cycle allows for nearly continuous read-out, such that dose fractionation has become ubiquitous as a means to record many-frame micrograph stacks in-place of traditional 2D images. The addition of a time-dimension, plus the large pixel counts of CMOS detectors, greatly increases both archival and data transfer requirements and associated costs to a laboratory. Many laboratories have a 1 Gbit/s Ethernet connection from their microscope to their computing center, which implies a data transfer rate of around 60–90 MB/s under typical conditions. If the microscope is run with automated data collection, such as SerialEM (Mastronade, 2005), then the so-called 'movie' may be 5–20 GB and may be saved every few minutes. In such a case, it may not be possible to transfer the data fast enough to keep up with collection. Costs for storing data on spinning (hard disk) storage, for example through the use of Google Cloud (Google Cloud Storage Pricing, accessed 2017), is typically US$100–200/TB/year. A cryo-TEM laboratory producing 200 TB of data per year is potentially faced with an annual data storage cost on the same order of magnitude as a post-doctoral fellow salary.

One approach whereby considerable archival savings may be realized is by decimation of the data from floating-point format to integer-format. Nominally, the analog-to-digital converted signal from the detector is typically output as a 16-bit integer. Due to data processing requirements, it is often necessary to convert the integer data to 32-bit floating point (decimal) format. The most common initial step that results in decimal data is the application of a gain reference, where the bias of the detector white values is removed. In-addition, conversion to floating-point is often inevitable due to operations such as sub-pixel shifting in drift correction (Li et al., 2013a, Grant and Grigorieff, 2015, McLeod et al., 2016, Zheng et al. 2017), or image filtration. If instead the micrographs are stored as 8-bit integers, with the gain reference (and potentially other operations) stored in meta-data, then a 4x reduction in storage and transfer requirements is realized. In this case, the gain reference and other bias corrections must be performed at the computing center, rather than using the software provided by the direct electron detector vendor. Since vendor gain normalization techniques are often proprietary and secret, there is a need for open-source equivalent solutions.

Further improvements in data reduction can be realized by modern high-speed lossless compression codes. Lossless compression methods operate on the basis of repeated patterns in the data. Nominally, purely-random numbers are incompressible. However counting electron data is Poisson distributed, such that its range of pixel histogram covers on only a limited range of values. In such a regime substantial compression ratios may be achieved. Therefore due to the repetition of intensity values, integer-format data can be compressed much more efficiently than gain-normalized floating-point data. Generally when comparing compression algorithms one is interested in the compression rate (in units of megabytes/second) and the compression ratio (in percent). Modern compression codecs such as Z-standard (github.com/facebook/zstd, accessed 03/2017) or LZ4 (github.com/lz4/lz4, accessed 03/2017) are designed for efficient multi-threaded operation on modern, parallel CPUs and can compress on the order of 1–2 GB/s/core, such that the time for read/write/transfer plus compression operations is greatly faster than when operating on uncompressed data.

We propose here combining decimation to 8-bit integer combined with lossless compression, with a robust post-acquisition application of the gain reference and detection and filtering of outlier (hot/dead) pixels. We propose an extension of the venerable MRC format, where *meta-compression* is implemented which implies the combination of loseless compression and loseless filtering to improve compressibility as well as execution with efficient blocked and multi-threaded processing. We propose a method to embed an unrestricted quantity of additional metadata in a footer of the MRC file, and compare JSON (ECMA-404, 2013) and Message Pack (msgpack.org, accessed 03/2017) encoders. The application of the gain reference by the detector vendors is often treated as a trade secret. We propose open-source outlier-pixel suppression and flat-field normalization algorithms.

## 2 The MRCZ format

The MRC format was introduced by Crowther et al. (Crowther et al., 1996) as an extension of the CCP4 format. It features a 1024-byte metadata header, followed by binary image data with provisons for 3-dimensions. The supported data types are byte (int8), short (int16), or single-precision floating-point (*float32*). The simplicity of the MRC format, and its ease of implementation, is a likely reason contributing to its popularity. However, the MRC format suffers from some drawbacks. First, there is no one standard format for MRC, in-spite of many efforts to define one (Cheng et al. 2015). Second, the header for storing metadata is only 1024-bytes long, so the quantity of meta-data that may be embedded is limited. Third, it cannot compress the data, so it is inefficient from an archival and transmission/distributed computing perspective.

An alternative public domain archival format for electron microscopy is HDF5. However, HDF5 is a “heavyweight” library consisting of ~350'000 of lines of code and 150-pages of specification (HDF Group, accessed 2017), which makes integration in existing projects difficult.

Here we introduce an evolution of the MRC format, MRCZ with additional functionalities that have become needed in the era of 'Big Data' in electron microscopy. We provide sample libraries for MRCZ in C/99 and also Python 2.7/3.5, as well as a command-line utility that may be used to compress/decompress MRC files so that legacy software can read the output. To facilitate the introduction of MRCZ into other software packages, we have kept the implementations as small as possible (currently *c-mrcz is* < 1000 lines of code).

The MRCZ library package leverages an open-source, meta-compression library, *blosc* (blocking, shuffle, compression), principally written by Francesc Alted and Valentin Haenel (Haenel, 2014, and Alted, accessed 03/2017). *Blosc* combines multi-threaded compression (currently six different codecs are available) with blocking, such that each operation fits in CPU cache (typically optimized to level 2 cache), and filter operations (namely *shuffle* and *bitshuffle)*. In testing on cryo-TEM data *blosc* achieved > 10 *GB/s* compression rates on a modern CPU, and superior compression ratios to codecs such as LZW (Welch, 1984) implemented in TIFF. The performance gain is sufficient such that loading a compressed image stack from disk and applying post-processing gain normalization and outlier pixel filtering to it is faster than loading an uncompressed but pre-processed floating-point result.

Here three compressors are compared for operation on cryo-EM data:

1. *lz4:*is the fastest compressor, with the worst compression ratio, making it ideal for live situations where distributing the data from a master computer is the priority.
2. *zStandard (zstd)*. achieves the highest compression ratio and has the fastest decompression, making it the best choice for archiving. On the lowest compression level it maintains good compression rates.
3. *Zlib:* is a very common library that has been accelerated by *blosc. Zlib* provides a valuable baseline for comparison, although *blosc* performance with *zlib* exceeds that of tools such as *pigz*

### 2.1 Blocked compression

Roughly around 2005, further increases in CPU clock-frequencies were slowed due to heat generation limitations. Further performance improvements where then realized by packing parallel arithmetic and logic cores per chip. Most common compression algorithms were designed before the era of parallel processing.

Operations in image processing are often relatively simple and executed on the full-frame consisting of many million elements. With the larger number of cores available on modern CPU, often program execution rate is limited not by processing power but the amount of memory bandwidth available to feed data to the cores. Typically fetching data from random-access memory (RAM) is an order of magnitude slower than the cache found on the CPU die. Therefore if the data can be cut into blocks that fit into the lower-level caches large speed improvements are often observed. Parallel algorithms can be made to work efficiently in the case where a computational task can be cut into blocks, and each block can be dispatched to an individual core, and run through an algorithm to completion, as illustrated in Fig. 1a. Parallel algorithms should also avoid branching instructions (e.g. conditional *if* statements), as modern processors request instructions from memory in-advance, and a wrong guess can leave the process idle waiting for memory. For example, *zStandard* also makes use a of a faster and more effective method for evaluating entropy, known as Asymmetric Numeral System (ANS), than classic compression algorithms. ANS significantly improves compression ratio in data with large degrees of randomness (Duda, 2013, and Duda et al., 2015).

**Figure 1:**
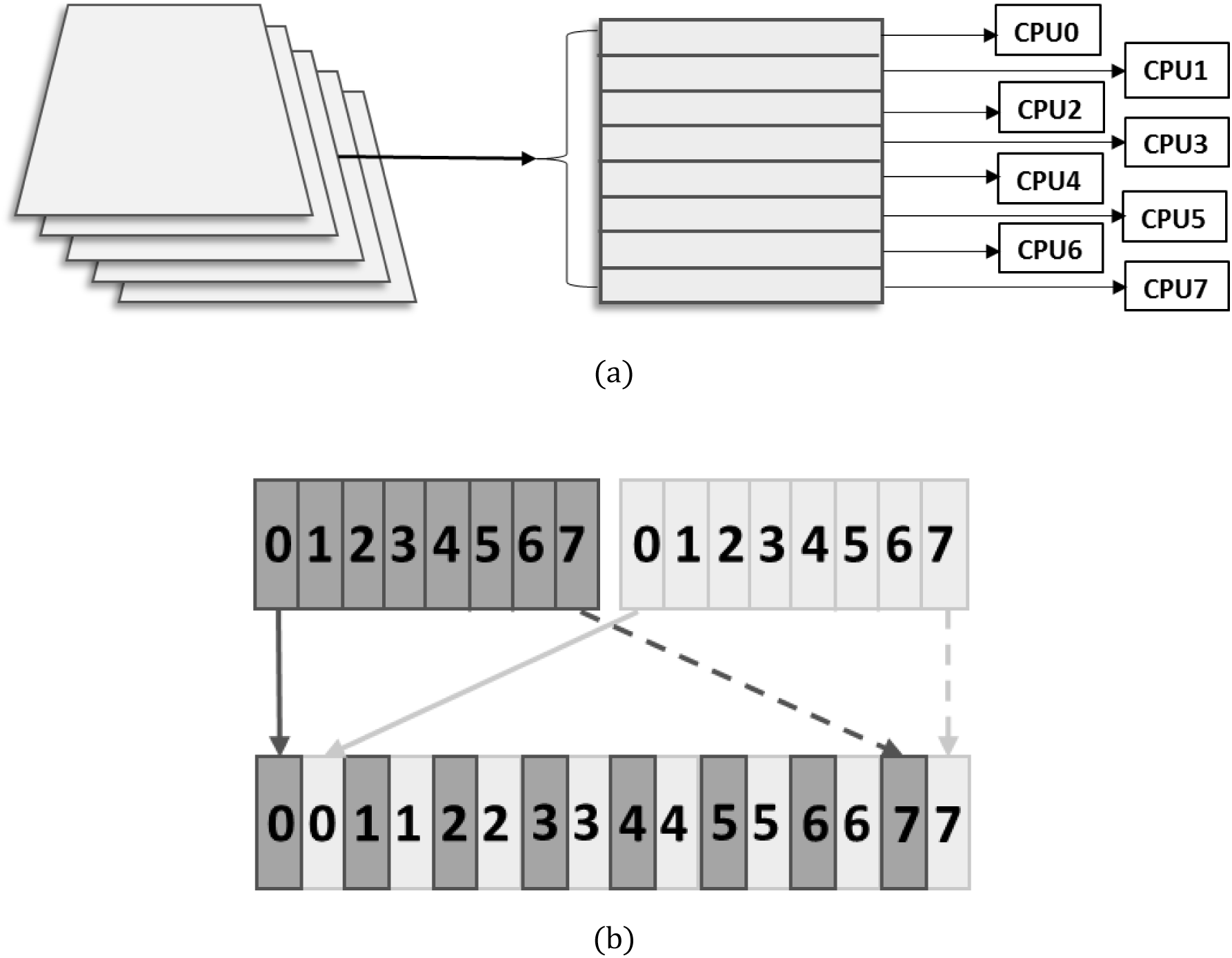
(a) Each MRC volume is chunked, such that each z-axis slice is compressed separately. Then in *blosc* each chunk is further sliced into blocks, which are then dispatched to individual CPU cores for compression. Decompression works in reverse. (b) Normally pixel values are stored in memory contiguously (top row). With bit-shuffling (on little endian systems) the most significant bits (**7** index) are stored adjacently, and similarly for the least-significant bits (**0** index). This improves compressibility and as a result both compression ratio and compression rate are improved.

In blocking strategies, data is conceptually separated into chunks and blocks, with chunks being senior to blocks. In the MRCZ format, each chunk is a single image frame (~16 million pixels for 4k detectors, or ~64 million for 8k), and each chunk is broken into numerous blocks, with a default block size of 1 MB. Such a block size provides a balanced trade-off between compression rate and the ratio between compressed and uncompressed data. For image stacks, the highest compression ratio would likely be in the time-axis, but this is the least convenient axis for chunking, as it would make retrieving individual frames or slices of frames impossible.

### 2.2 Bit-decimation by shuffling

Some direct electron detectors, such as the K2 Summit (Gatan, Pleasanton, CA) cycle fast enough to count individual electrons during image acquisition. Counted electron images are often dose fractionated whereby each time slice has mean dose < 4 *e^−^/pix/frame*. This implies that even a single byte *(uint8)* to store each pixel is too large of a data container, as it can hold data values up to 255. David Mastronde implemented in *SerialEM* and *IMOD* (Mastronade, 2004) a new data type for MRC that incorporates a decimation step where each pixel is packed into 4-bits, leading to maximum per-pixel values of 16 before clipping occurs, thereby providing an effective compression ratio of 2.0 compared to *uint8*. The disadvantage of 4-bit packed data is that it is not a hardware data type, such that two pixels are actually packed into an 8-bit integer. Whenever the data is loaded into memory for processing, it must be unpacked with bit-shifting operations, which is computationally not free. There is also the risk of intensity-value clipping.

*Blosc* optionally makes use of a filter step, of which there are two currently implemented, *shuffle* and *bitshuffle. Shuffle* re-arranges each pixel by its most significant byte to least, whereas *bitshuffle* performs the same task on a bit-level, illustrated in Fig. 1b. The shuffle-style filters are highly efficient when the underlying data has a narrow histogram, such that the most-significant digit in a pixel has more commonality with other pixels’ most significant digit than its own least-significant digit. For example, if an image is saved as *uint8* type contains mostly zeros in its most significant digits, they will be bit-shuffled into a long-series of zeros, which is trivially compressible. As such, *bitshuffle* effectively performs optimized data decimation without any risk of clipping values. Shuffling is also effective for floating-point compression, as the sign bit and the exponent are compressible whereas the mantissa usually does not contain repeated values and therefore it is not especially compressible. The mantissa can be made more compressible by rounding to some significant bits, for example the nearest 0.001 of an electron, but this generates round-off error.

### 2.3 Benchmarks

Benchmarks for synthetic random Poisson data were conducted for images covering a range of electron dose levels consisting of [0.1, 0.25, 0.5, 1.0, 1.5, 2.0, 4.0] electron counts/pixel. The free parameters examined consist of: compression codec, blocksize, threads, and compression level were all evaluated. Here the term ‘compression level’ refers to the degree of computing effort the algorithm will use to achieve higher compression ratios. The machine specification for benchmark results is as follows:

Two Intel^®^ Xeon^®^ E5-2680 v3 CPUs operating with Hyperthreading^®^ and Turboboost^®^:

- No. of physical cores: 2 × 12
- Average clock rate: 2.9 GHz (spec: 2.4 GHz)
- L1 cache size: 32 KB per core
- L2 cache size: 256 KB per core
- L3 cache size: 30720 KB per processor

The size of the L2 cache generally has a large impact on the compression rate as a function of the blocksize used by *blosc*. For file I/O the RAID0 hard drive used was benchmarked to have a read/write rate of ~300 MB/s, which is comparable to parallel-file systems in general use in cluster environments.

Example benchmarks on cryo-TEM image stacks are shown in Table 1 for a variety of *blosc* libraries as well as external compression tools. *Uint4* refers to the *SerialEM* practice of interlaced packing of two pixels into a single-byte. JPEG2000 and *uint4were* not multi-threaded; all other operations used 48 threads. *Pigz*, *lbzip2* and *pxz*are command-line utilities and hence include a read and write. Indicated times are averages over 20 read/writes. To achieve repeatable result, the disk was flushed between each operation, with the Linux command:
`echo 3 | sudo tee /proc/sys/vm/drop_caches`

**Table 1:** Comparison of read/write times for 60 × 3838 × 3710 cryo-TEM image stacks.

| Codec<br>/ data type<br>/compression level | Compressed Size<br>(MB) | Compression<br>Ratio | Compression-Write Time<br>(s) | Decompression-Read Time<br>(s) |
| --- | --- | --- | --- | --- |
| None / int8 | 854 | 1.00 | 3.40 | 3.21 |
| uint4 | 427 | 0.50 | 2.14 | 6.05 |
| blosc-lz4 / int8 / 9 | 340 | 0.40 | 0.50 | 0.96 |
| blosc-zstd / int8 / 1 | 320 | 0.37 | 0.76 | 1.10 |
| blosc-zstd / int8 / 5 | 319 | 0.37 | 0.86 | 1.09 |
| JPEG2000 / uint8 | 317 | 0.37 | 106.8 | N/A |
| pigz / int8 / 1 | 367 | 0.43 | 0.86 | 4.74 |
| lbzip2 / int8 / 9 | 314 | 0.37 | 3.17 | 2.23 |
| pxz / int8 / 6 | 305 | 0.36 | 47.5 | 31.8 |

The gains in compression ratio by using more expensive algorithm such as Burrows-Wheeler (bzip2) or LZMA2 (xz) are quite minimal with cryo-TEM data, likely due to the high degree of underlying randomness (or entropy). *Lbzip2* is the clear winner among command-line compression tools, as it still is faster than reading or writing uncompressed data and achieves the second-best compression ratio. *Blosc* accelerates the read/write by a factor of 3–6x over that of the uncompressed data.

Figures for benchmark results are shown in Figure 2. Best compression ratio as a function of dose rate is shown in Figure 2a. An important consequence of compressing Poisson-like data is that compression ratios increase substantially with sparseness. I.e. compressed size scales sub-linearly with decreasing dose fractions. For example, a cryo-tomography projection of 10 frames of 4k × 4k data recorded at a dose rate of 0.1 *e^−^/pix/frame* would have a compressed size of 13 MB, compared to 670 MB for its uncompressed, gain-normalized image stack. Such compression therefore enables finer-dose fractionation for advanced drift correction algorithms without imposing onerous storage requirements.

**Figure 2:**
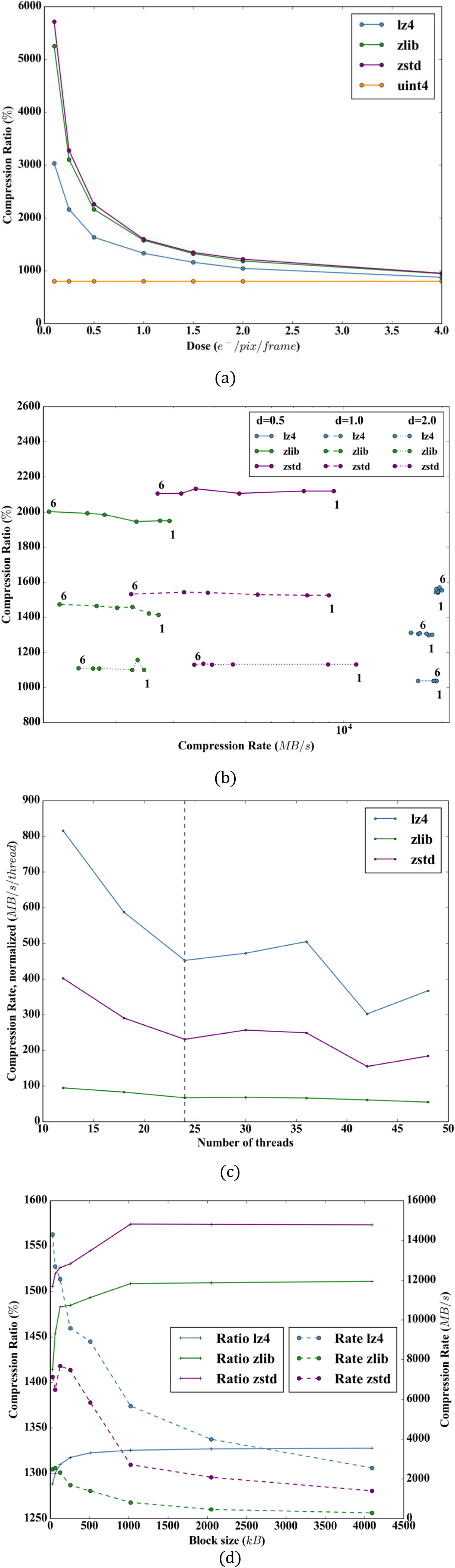
Performance for various compression codecs found in *blosc*. (a) The dependence of compression ratio varies strongly with the dose. Here *zstd has* the best compression ratio. (b) The dependence of the compression level on the compression ratio is mild, such that for *zstd* and *zlib* typically 1 is used. (c) Scaling on the compression rate with the number of parallel computing threads employed. The machine used has 2×12 physical cores, indicated with the dashed line. The area to the right of the dashed line indicates the region in which Intel Hyperthreading^®^ is active. (d) Dependence of the compression ratio and rate on the blocksize used, which is the most critical parameter examined. Typically a blocksize scaled to fit into L2 cache (256 *kB*) is optimal for speed, but the compression ratio benefits from a larger blocksize (≥ 512 *kB*).

With regards to compression level, shown in Figure 2b, which is a reflection on the effort level of the compressor, generally *zlib* saturates at 4–5, whereas *zstd* saturates at 2–3, and *lz4* sees little disadvantage to running at its highest compression level. Good compromises for processing are compression level 1 for *zstd* and *zlib* and for archiving 3 for *zstd* and *5* for *zlib. Lz4* can operate with compression levels of 9 for real-time applications but it is not as suitable for archiving due to the lower compression ratios. The *bitshuffle* filter is important in this situation and contributes heavily to the quickest compression level of 1 being the best compromise between rate and ratio for *zstd*, in that it uses *a priori* knowledge about the structure of the pixel values to pre-align the data into its most compressible order.

In *blosc* the scaling with threads is roughly 1/0.7N_*threads*_ for *N_threads_* ≥ 2, up to the number of physical cores. When hyper-threading is enabled an oversubscription of approximately *N_threads_* ≈ 1.5 *N_cores_* gives the highest absolute compression rate.

Cache sizes are important in that they impose thresholds on data sizes, shown in Fig. 1d. *blosc* chops the data into blocks, and *MRCZ* cuts a volume into single z-axis slices called *chunks*. For example a 4 k × 4k image chunk may be cut into 64 separate 256 kB blocks. If the block size fits into the L2 cache (≤ 256 *kB*) then compression rate advantage is expected, and this is evident. However, testing on simulated Poisson data shows that larger blocks (which result in larger dictionaries in the compression algorithm) achieve a higher compression ratio. Similarly for chunking, if the chunk size is less than the L3 cache (≤ 30 *MB)* then only one memory call is sufficient for the entire chunk.

The optimal block size for compression ratio is expected to be when each block holds one significant bit each. So for 4k × 4k × 8-bit images the ideal block size would be 16/8 *=* 2 *MB*, whereas the saturation in compression ratio actually appears at 1 MB.

### 2.4 Enabling Electron Counting in Remote Computers with Image Compression

The K2 Summit direct electron detector is able to count single electrons by sampling fast enough (400 Hz) such that two electrons are unlikely to impact the same pixel in the same sampling period. The raw data is transferred to a field-programmable gate array (FPGA) processor, which conducts thresholding operations in real-time to count and optionally estimate electron impact points with subpixel precision by center-of-mass determination. Such a device is a remarkable achievement, but the user does not have access to the raw data largely due to the enormous data rate (5.7 GB/s) at which the images are recorded. This is unfortunate as it does not permit experimentation with alternative subpixel detection algorithms that might better localize the impact of the primary electron in the detector layer.

However the multi-threaded meta-compression discussed above may be able to alleviate the data flow problem. A Gatan K2 detector samples at 3838 × 3710 × 400 Hz, or 5.7 GB/s, and the FEI Falcon 3 at 4096 × 4096 × 100 Hz = 1.7 GB/s. Both throughputs are less than that demonstrated above for *blosc*. A master node could, using *zstd* (or falling that, *lz4*) as a compression codec, compress the raw data from a K2 on-the-fly and dispatch it to worker computers for counting, thus enabling counting without a FPGA or similar hardware counting solution. The compression ratio achievable would depend heavily on thresholding of low intensity values but could be in the range of 50:1.

## 3 Extended Metadata in MRCZ

The MRC2014 format allows for the potential of an extended header that is longer than the standard 1024 bytes. The disadvantage of this approach is that is it not backward compatible with previous MRC file formats where the programmer could safely assume the data begins at byte 1024. In order to preserve backward compatibility with the original MRC specification we propose that instead extra meta-data be added as a footer to the MRC data. This is backward compatible (when using uncompressed data) with all tested MRC using software (Relion, Frealign, SerialEM, Digital Micrograph, EMAN2) as they simply ignore the presence of the extra meta-data.

File formats require complex, nested metadata to be encoded into a stream of bytes. The conversion of metadata to bytes is called serialization. In order to achieve a high level of portability into the future for the metadata, we advise use of a well-established serialization standard. Here two serialization standards are compared, JSON (JavaScript Object Notation), which is the most ubiquitous serialization method in the world, and Message Pack (msgpack.org and pypi.python.org/pypi/msgpack-python). a binary serialization tool with a similar language structure to JSON. Libraries are available for both for many different programming languages, with the exception of Matlab and Fortran for Message Pack. Here two high-speed JSON encoders available for C and Python are profiled, RapidJSON (rapidison.org and pypi.python.org/pypi/pyrapidison) and UltraJSON (github.com/esnme/uison4c and pypi.python.org/pypi/ujson). The three serialization methods were tested on a sample of 25 MB of complicated JSON data. All three methods produce more-or-less similar results in terms of read/write times to disk, as shown in Fig. 2a. Compression does not speed nor slow read/write times, except for Message Pack read rates. *Lz4* reduces the data size on disk to roughly one-half, and *zstd* to roughly one-third, of the uncompressed size. Message Pack was tested with Unicode-encoding enabled to make it equivalent to the JSON encoders, which slows its read/write time by ~20 %.

## 4 Stochastic outlier pixel suppression algorithm

With direct electron detectors the active layer and CMOS electronics suffer constant radiation damage during data acquisition. As such, outlier pixels apparently are far more common than in traditional scintillator coupled CCD detectors, and they also appear to be far more dynamic, appearing and disappearing from image to image. Outlier pixels are effectively Dirac functions in real-space, which makes them impractical to filter in reciprocal-space as they span all frequencies. Outlier pixels appear as statistically anomalous “fat tails” to the expected distribution of a compound Poisson distribution (which can be well approximated by a normal distribution). They can be dead pixels, with low values, which are relatively rare, or hot pixels, which with direct electron detectors are far more likely. Typically around 0.5 % of the pixels on a Gatan K2 are found to be outliers using the approach detailed below. The problem is illustrated schematically in Fig. 3. A synthetic particle has suffered motion blur due to drift, but the outlier pixels are stable in position over the course of the dose fractionation. After drift correction, ideally the particle motion is compensated for but the outlier pixels have not.

**Figure 3:**
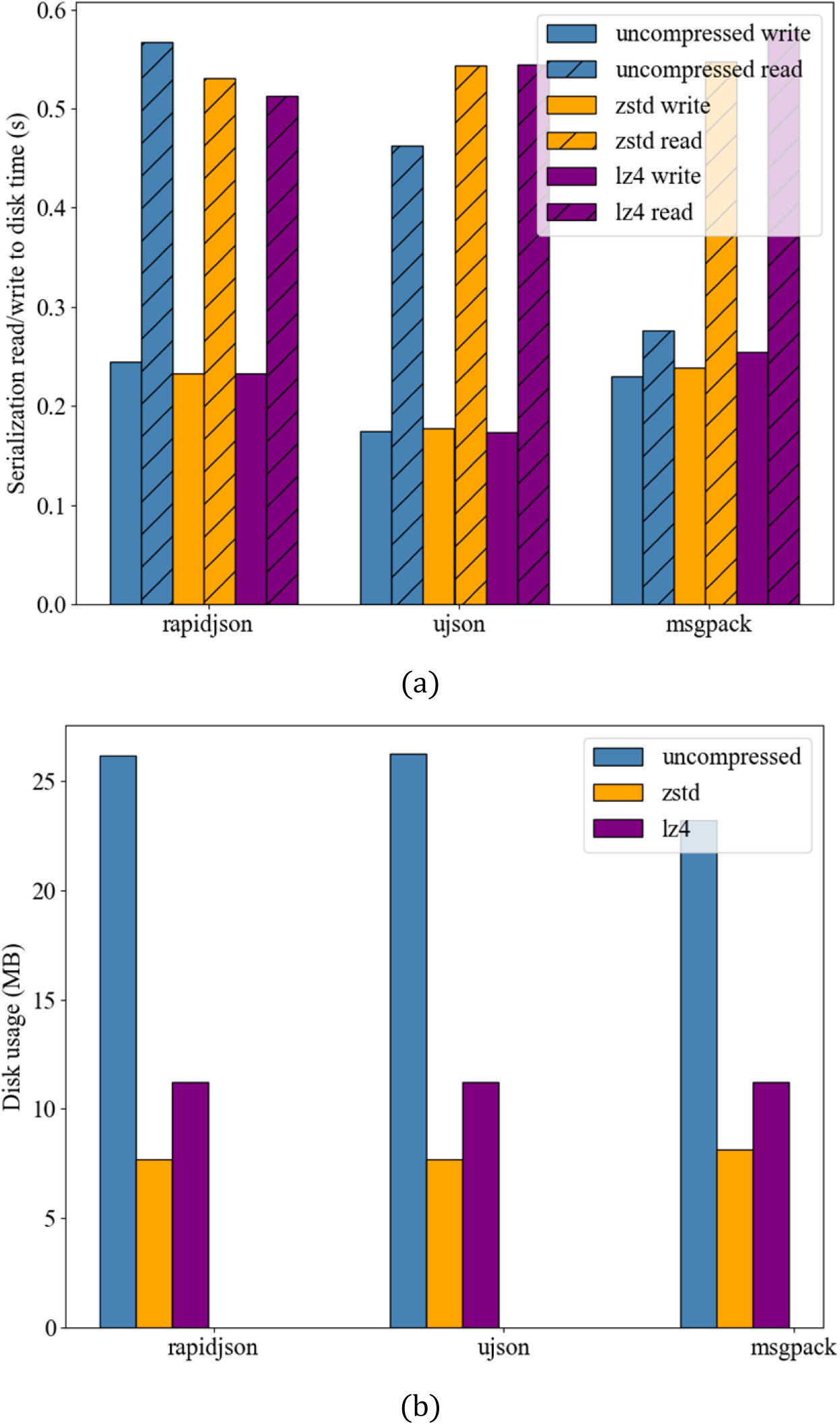
Performance evaluation of different serialization methods for meta-data paired with compression. (a) Read and write times for the profiled serialization methods on a sample of 25 MB of JSON-like text metadata data, when used with and without blosc compression. (b) Size on disk of the metadata.

Common approaches to detecting outlier pixels find either those that are spatially anomalous, by measuring the standard deviation of a pixel compared to the neighbors or the ensemble average, or temporarily anomalous, by measuring the standard deviation in time. Here the approach is to examine for spatial anomalies by detecting pixels that deviate strongly from a normal distribution in the image histogram. Computational time for the outlier pixel removal algorithm is ~2–3 s for a 50 × 4096 × 4096 image stack. The algorithm is described as conceptual steps:

Step 1: Calculate a histogram of the stack summed along the time axis with electron count-centered bins over the range (−8σ, 8σ). The unaligned sum is used as the outlier pixels will reinforce their intensities whereas the specimen contrast will be somewhat blurred by the motion during the acquisition. The cumulative summation of the histogram represents the estimate for the experimental cumulative distribution function CDF_*hist*_, and is typically normal for cryo-TEM micrographs. Samples with visible carbon may have a bi-normal histogram. All fit estimates are conducted on the cumulative rather than density function because, due to the numerical integration, it is more robust against noise.
Step 2: Fit the histogram cumulative distribution function with a normal CDF:

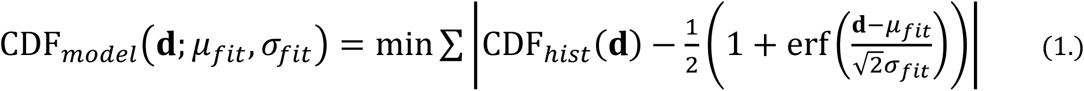

where **d** is the dose axis of the histogram, *μ_fit_* is the *σ_fit_* mean, and the best-fit standard deviation and erf the error function. In this work functional minimization was performed with the L-BFGS-B functional minimization algorithm but a least squares approach would likely suffice.
Step 3: Typically outlier pixels are described in terms of *sigma values*, or multiples of the standard deviation: transform the experiment CDF_*hist*_ from counts to terms of standard deviations (i.e. σ- values),

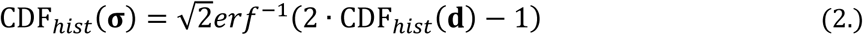

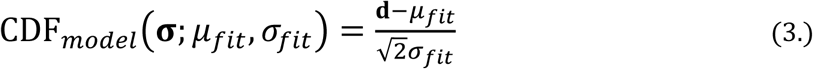 And then the residual difference *r* is

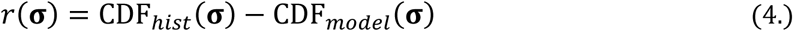 An example residual is shown in Fig. 4 as the orange line.

**Figure 4:**
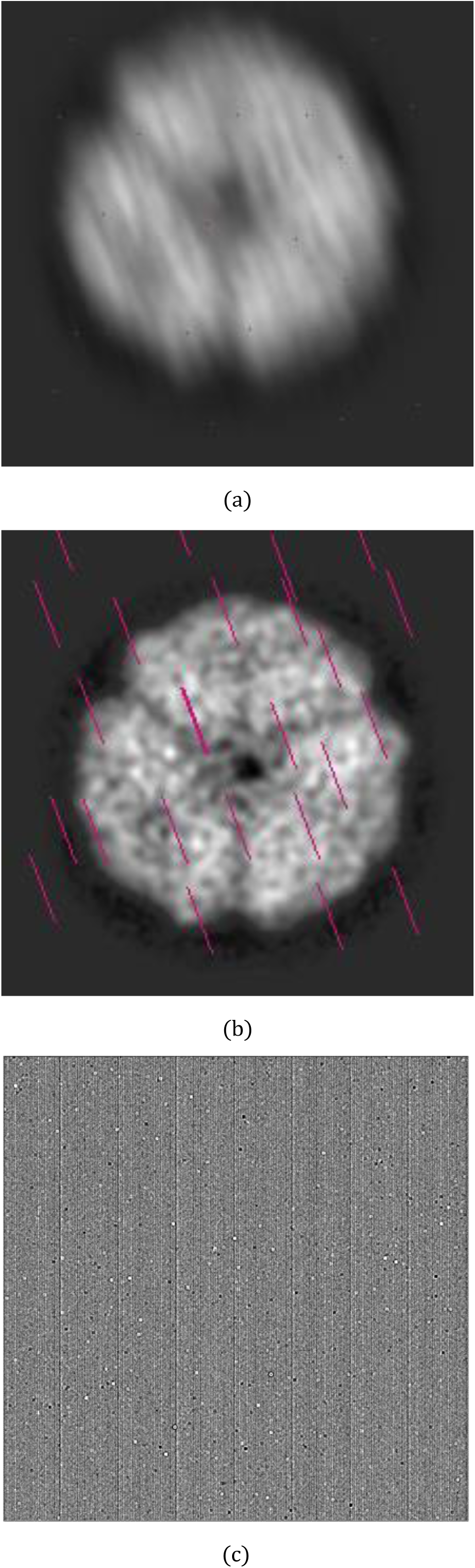
Scheme illustrating the effect of outlier pixels when drift correction is applied. (a) Representation of an unaligned average of a protein particle. Outlier pixels are represented by magenta dots and are stationary. (b) After drift correction, the aligned average shows a crisp projection of the particle but the outlier pixels are dragged across the frame, resulting in the shown magenta lines. The drift tracks contribute correlated noise to the aligned particle image. (c) An example 512 × 512 box from a K2 super-resolution gain reference shows approximately 200 outlier pixels. The CMOS pattern is visible as vertical lines in this case.

**Figure 5:**
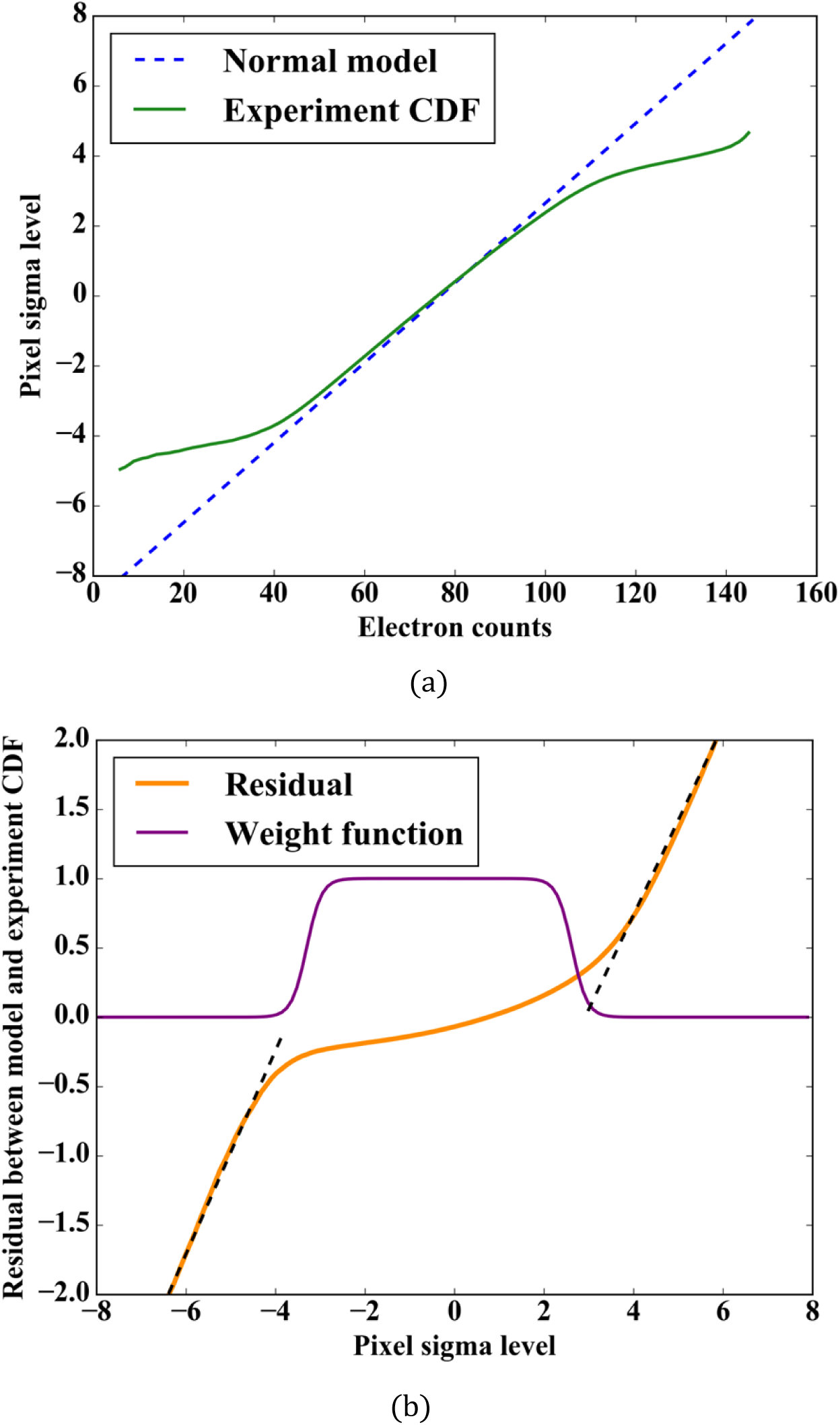
Outlier detection is performed with the cumulative histogram (i.e. cumulative distribution function) of the stack average. (a) The cumulative histogram of the image (green line), after intensity values are transformed to sigma values, by dividing by the Poisson standard distribution, 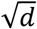, compared to the normal distribution model (blue dashed line). The ‘fat’ outlier tails onset at sigma values of ~(–4.0,3.25). (b)The residual difference between the model and experiment (orange line). The zero-intercepts of the best-fits to the fat tails (black dashed lines) are used as the thresholds for dark and hot pixels. A logistical weighting function (purple line) is used to determine the proportion of the original pixel value that is retained by the filter.

**Figure 6:**
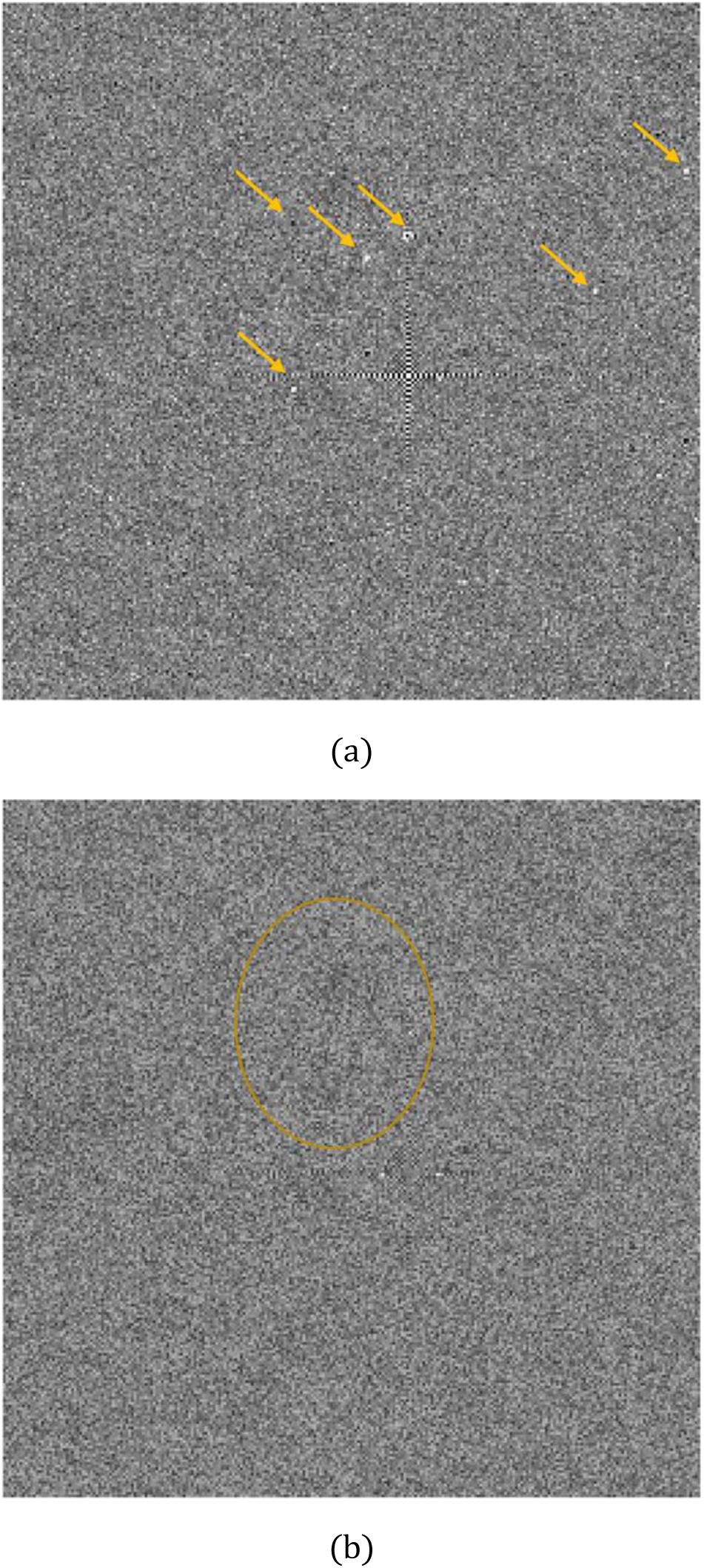
Example of a recorded cryo-EM image of a vitrified 160 kDa protein particle containing several examples of outlier artifacts. (a) Pre- and (b) Post-outlier pixel filter results for a 256 × 256 patch. Arrows in (a) indicate locations of outlier clusters, and the circle in (b) shows the 160 kDa particle, demonstrating that the contrast of outlier pixels can be quite strong relative to the contrast of small protein particles.

**Video 1 (available on-line):**
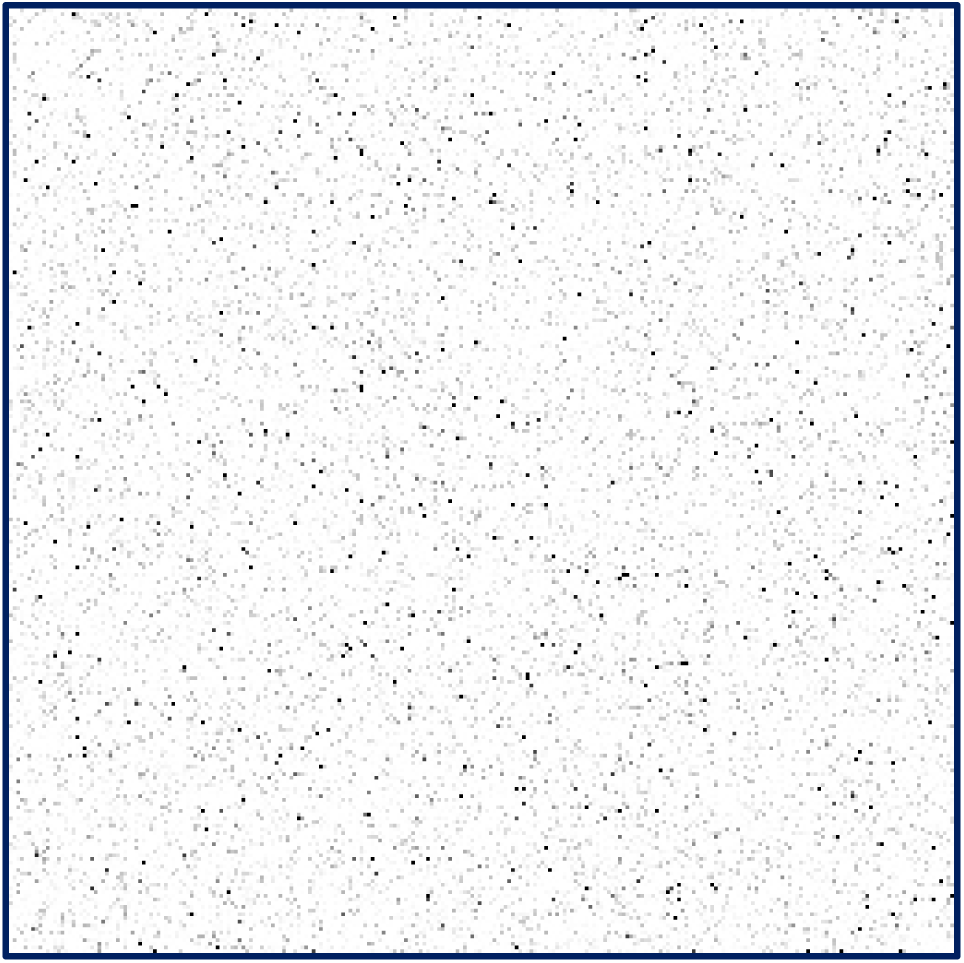
The distribution of outlier pixels is tracked over ~6 hours of single particle data acquisition. The still frame shows the average proportion of outliers, which shows that outlier distribution is more-or-less random.
Step 4: Fit the left- and right-side large residuals with linear polynomials to find the respective zero residual intercepts for dark pixels, *t_dark_*, which are outliers on the black end of the histogram, and hot pixels, *t_hot_*, which are outliers on the white end of the histogram.

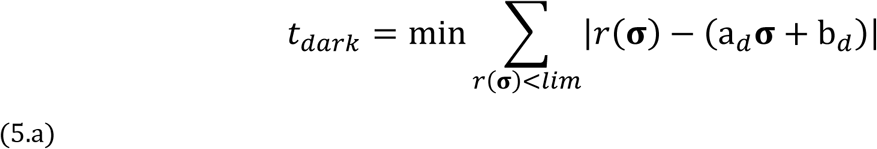

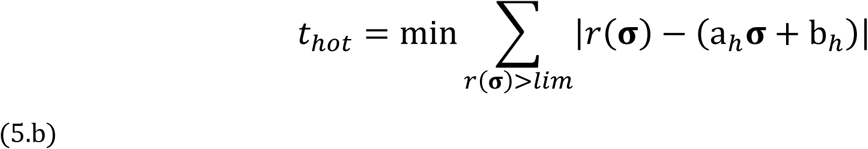 The fit is only conducted over the σ-range where the residual magnitude |*r*(**σ**) | is greater than some limit, typically *σ =* 0.5. From these thresholds a logistic weighting function *w* is constructed,

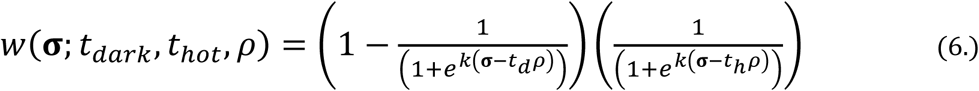

where *k* is the filter order and defaults to 6.0, and *ρ* is a relaxation of the threshold for which a typical value is 0.95.
Step 5: Compute the outlier pixel mask. Two approaches are used to fill in outlier pixels. *Singletons* are outlier pixels that have no outlier neighbors within a two pixel radius. In this case, the estimated point-spread function of the detector is used to estimate the expectation value for the outlier pixel; the estimated fill-in values for the Gatan K2 are shown in Table 1 (negative shifts are not shown as the function is symmetric). Non-singleton *neighborly pixels*, i.e. those with outlier neighbors within a two pixel radius, can constitute as high as 15 – 25 % of the total outlier pixels in normal mode and > 66 % in super-resolution mode and must be handled differently. Here the mean (not including any outlier pixels) of a 5 × 5 neighborhood about each outlier replaces the outlier pixel value.

**Table 1:** Gatan K2 point-spread function as a function of neighbor adjacency.

| Y / X | 0 pix shift | 1 pix shift | 2 pix shift |
| --- | --- | --- | --- |
| 0 pix shift | 0 | 0.173 | 0.03 |
| 1 pix shift | 0.173 | 0.084 | 0.02 |
| 2 pix shift | 0.03 | 0.02 | 0.007 |

### 4.1 Effect of outlier suppression on correlated noise

In order to evaluate the capacity of the algorithm to reduce correlated noise, 500 dose fractionated image stacks were acquired, each consisting of 100 frames each with an exposure time of 300 *ms/frame*, and a dose rate of 4 *e^−^/pix/s*. The detector was annealed prior to data acquisition and then a fresh gain reference acquired, with a gain factor of 100, which was applied to each image in the set. As shown in Fig. 7(a) the number of hot pixels rises over the course of 500 image stacks, from approximately 5’500 to 12’300, representing > four hours of continuous imaging. The dead pixel count is more stable, rising from approximately 160 to 180. A similar experiment was conducted with the detector in a mature-state, at a higher dose rate (7 *e^−^/phys. pjx/s*), and in super-resolution mode. Here the proportion of outlier pixels was approximately 10x higher, as shown in Fig. 7(b), but the number of outliers does not appear to increase with cumulative dose.

**Figure 7:**
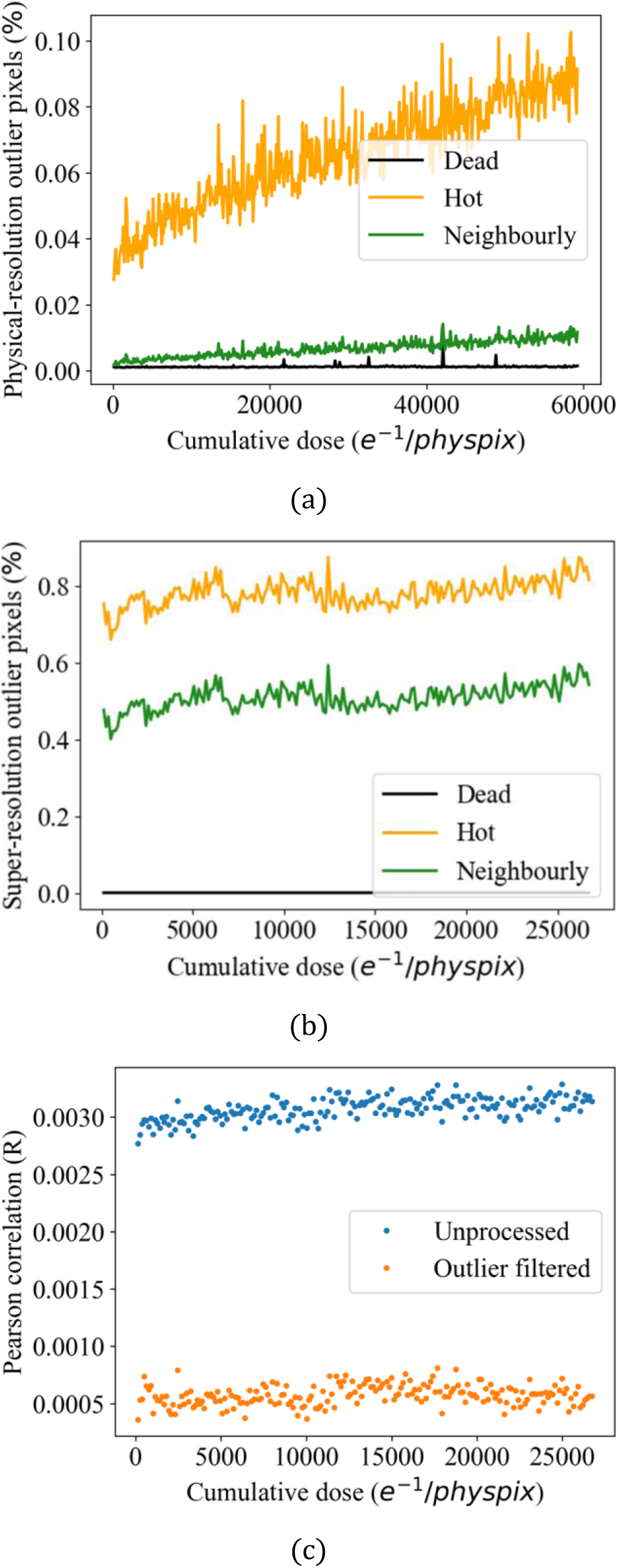
Analysis of the density of outlier pixels as a function of cumulative electron dose for two cases. (a) For normal resolution mode of a K2 Summit, plotting the percentage of outlier pixels versus cumulative dose starting from an annealed-state at a dose rate of 4 *e^−^/pix/s* shows the rise in the number of hot pixels over approximately four hours of operation. (b) An analysis of the detector from a mature-state in super-resolution mode at a dose rate of 7 *e^−^/pix/s*. Here because each physical pixel contributes to four super-resolution pixels the proportion of outlier pixels is much larger. The proportion of outlier pixels appears to have saturated and does not increase significantly with dose. (c) The amount of correlated noise among super-resolution frames in the mature-state is reduced by approximately 5-fold after the outlier filter is applied.

**Figure 8:**
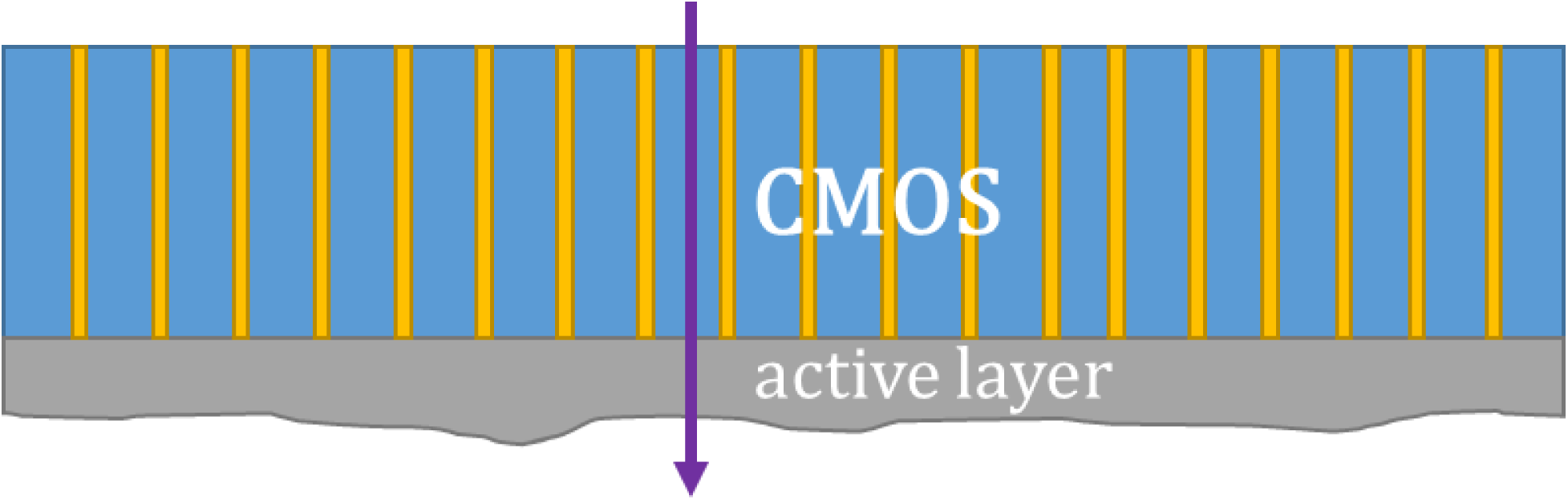
Schematic depiction of a K2 sensor. The incoming electron passes through the CMOS electronics first before impacting the active layer. The active layer is back-thinned, often some thickness variation due to etching is present (exaggerated here). Therefore the image on the detector is modified both by the patterns in the CMOS electronics, and the etching of the active layer, each with different dose-rate scaling.

To assess the degree of stationary correlated noise the Pearson correlation coefficient was employed, which can be calculate from the covariance of two images. For two images *X* and *Y*, the Pearson correlation *R* is computed as,

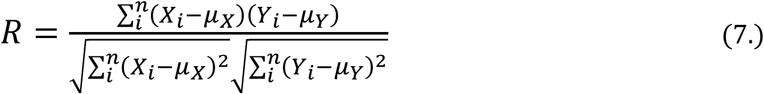

As shown in Fig. 7(c) the outlier filter reduces the correlated noise among frames inside an individual stack by approximately 550 %. The major source of residual correlated noise after outlier pixel suppression is thought to be caused by CMOS read-out lines.

## 5 Image normalization

Image normalization (also known as ‘flat-field correction’ or ‘gain reference multiplication’) is a standard method to suppress detector artifacts in digital imaging. Image normalization is particularly important when performing dose-fractionated image correction, as the fixed-pattern noise can often significantly displace template-matching methods from the true image drift. Historically, image normalization for CCD detectors consisted of subtraction of a dark reference image followed by multiplication by the gain reference,

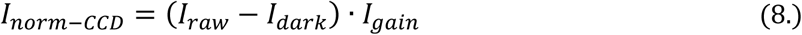

where *I_gain_* was typically the reciprocal of a ‘flat-field’ average of images obtained by illuminating the detector without a specimen in the field-of-view. The reciprocal is pre-screened for zeros, which cause divide-by-zero errors. With the Gatan K2 Summit, the normalization is believed to be applied in hardware, then thresholding is applied to count electrons. After accumulation electrons sampled at 400 Hz into a dose-fractionated frame, second gain reference is applied in software. It is likely that the second, software gain reference is more than a simple multiplication, as the gain reference changes significantly with dose rate, and there is also a need to further suppress features such as CMOS read-out lines.

An alternative approach to image normalization is to perform correction after acquiring a large volume of data, i.e. an *a posteriori* approach (Afanasyev, 2015). *A posteriori* approaches are inherently less convenient than *a priori* approaches because they require the research to wait but potentially can improve on the reduction in correlated noise. In section 5.1, a first effort on an equivalent-to-vendor *a priori* gain normalization is presented, whereby the gain reference is scaled by the dose rate with a gamma-factor. In section 5.2 the *a posteriori* approach is examined and a methodology suitable for pipeline processing is presented.

### 5.1 Gain variation with dose rate

Here *a priori* approaches to improve on Eqn. 8 are presented. With direct electron detectors it is apparent that the gain normalization is a function of the dose rate. As such, an additional gamma correction has been evaluated when applied to the gain reference. For a dose rate **δ**, the proposed image normalization equation is,

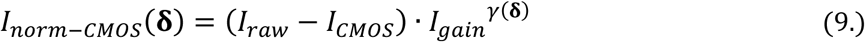

where *I_gain_* is the vendor provided gain reference, *γ* is a gamma correction to account for the active layer etching pattern, and *I_CMOS_* is a subtraction term designed to remove the overlain pattern of the CMOS electronics. A number of approaches were examined to find a suitable approach to compute *1_CMOS_ a priori*, such as Fourier filtering to generate electronic seam maps, but have been found to be unsatisfactory compared to the limited *a posteriori* approach in Section 5.2.

The best gamma for each dose rate ***δ*** is determined by minimizing the root-mean-square difference

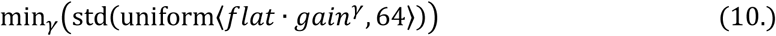

where uniform<…, 64> is the application of a uniform filter over a 64 × 64 pixel patch, to minimize high-frequency components, std(…) computes the standard deviation over the entire field-of-view after filtering, and mm_*γ*_ (…) is a functional minimization with *γ* as the free parameter. Example data were collected over a range of dose rates (0.25,9) *e^−^/pix/s*. The gamma normalization in Fig. 9 for a dose rate *δ* is fit to a power-linear function,

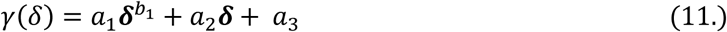

**Figure 9:**
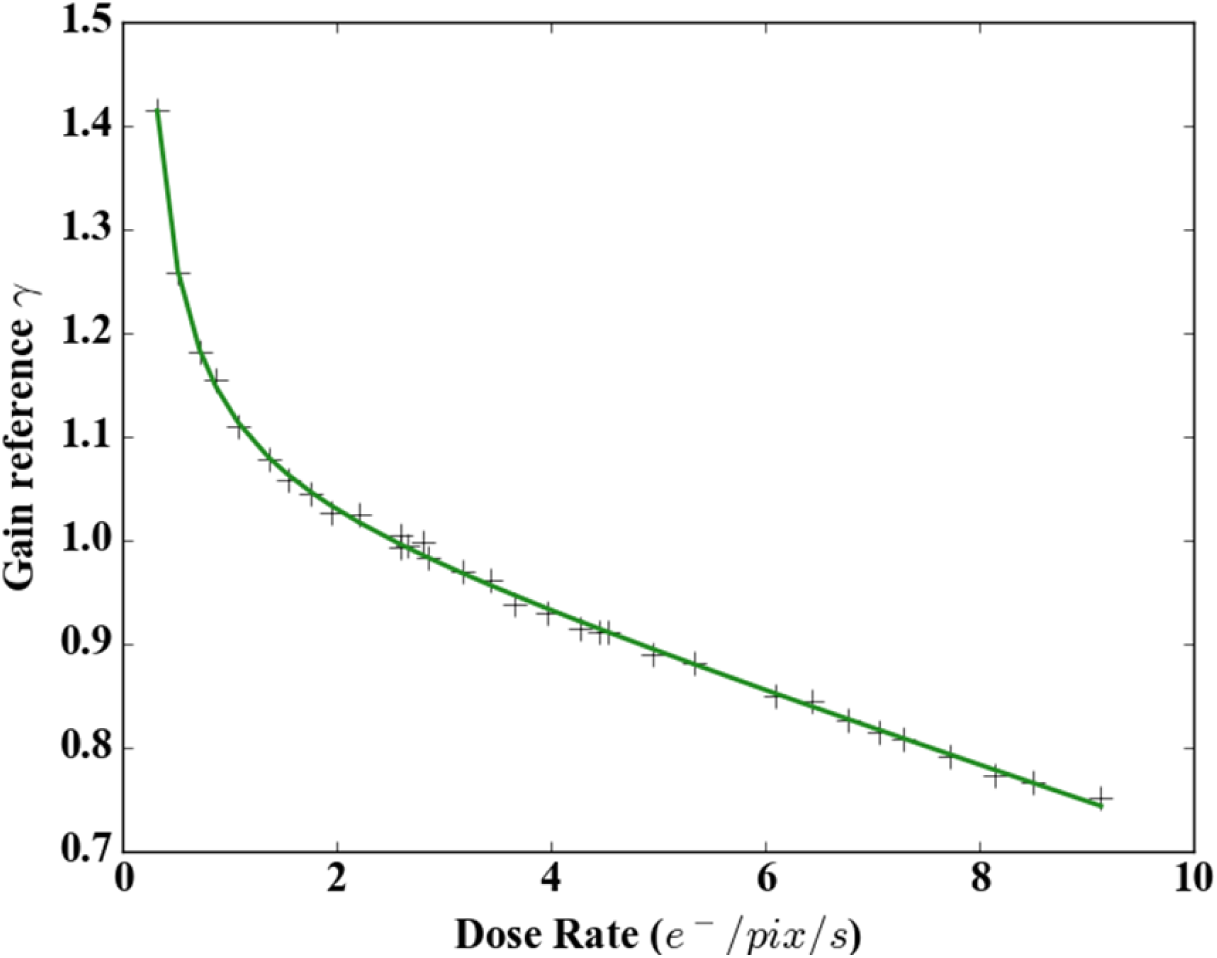
Plot of optimized gain reference gamma-factor versus dose rate. Green line is the best-fit of a power plus linear function to the data (crosses).

where *a_n_* and *b_n_* are free best-fit parameters (*a_1_* = 0.12, *a_1_ =* −0.0336, *a_3_ =* 1.04, *b_1_ =* −1.04). The gamma normalization does effectively remove the variation due to the etching of the active layer, as shown in Fig. 10, but it is ineffective in removing the CMOS electronics pattern.

**Figure 10:**
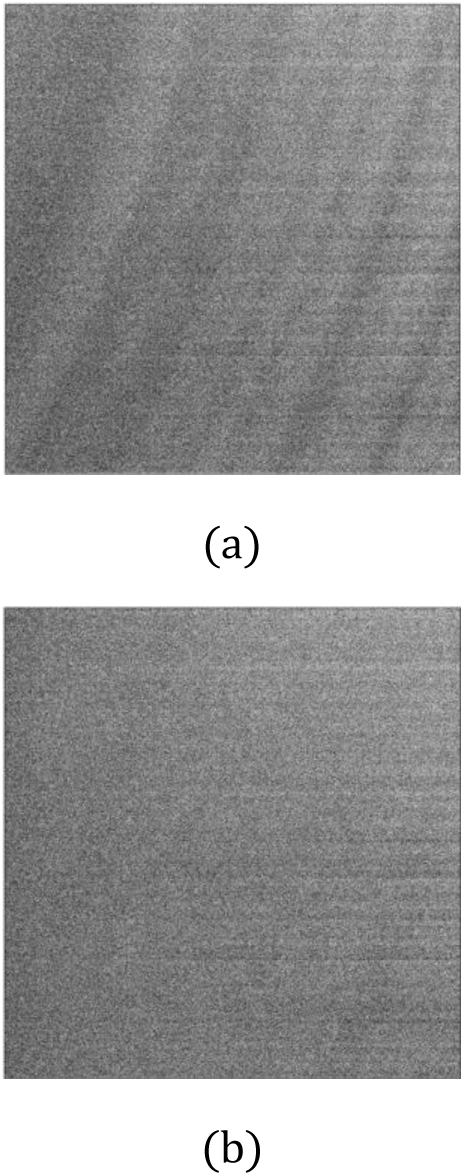
Example vacuum reference at a dose rate of 7 *e^−^/pix/s* of (a) typical-normalization as in Eqn. 7 and (b) gamma-normalization with *γ =* 0.85 in Eqn. 8. Images have been low-pass filtered to better show the etching pattern (diagonal sweeps). The CMOS read-out electronics are still evident in (b) as horizontal-lines on the right-side.

### 5.2 Knock-out Moving Average (KOMA) Normalization

In the *a posteriori* approach it is desirable to assess how many stacks must be averaged in order to generate an efficient normalization term *I_CMOS_* for Eqn. 9. In vacuum reference images, the correlation between stack averages drops rapidly with increasing separation in the dose (or time) axis as the detector is dynamically annealed in-situ by the electron beam. A series of 30 s dose-fractionated stacked were acquired over a period of four hours on an FEI Polara operated at 300 keV equipped with a Gatan K2 Summit detector, constituting over 50'000 electrons per pixel total dose, with the correlation of the stack sums (blue points) to the first stack sum shown in Fig. 11. The fit to the vacuum correlation with cumulative dose (gold line) shown in Fig. 11 is,

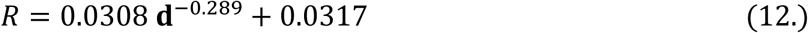

**Figure 11:**
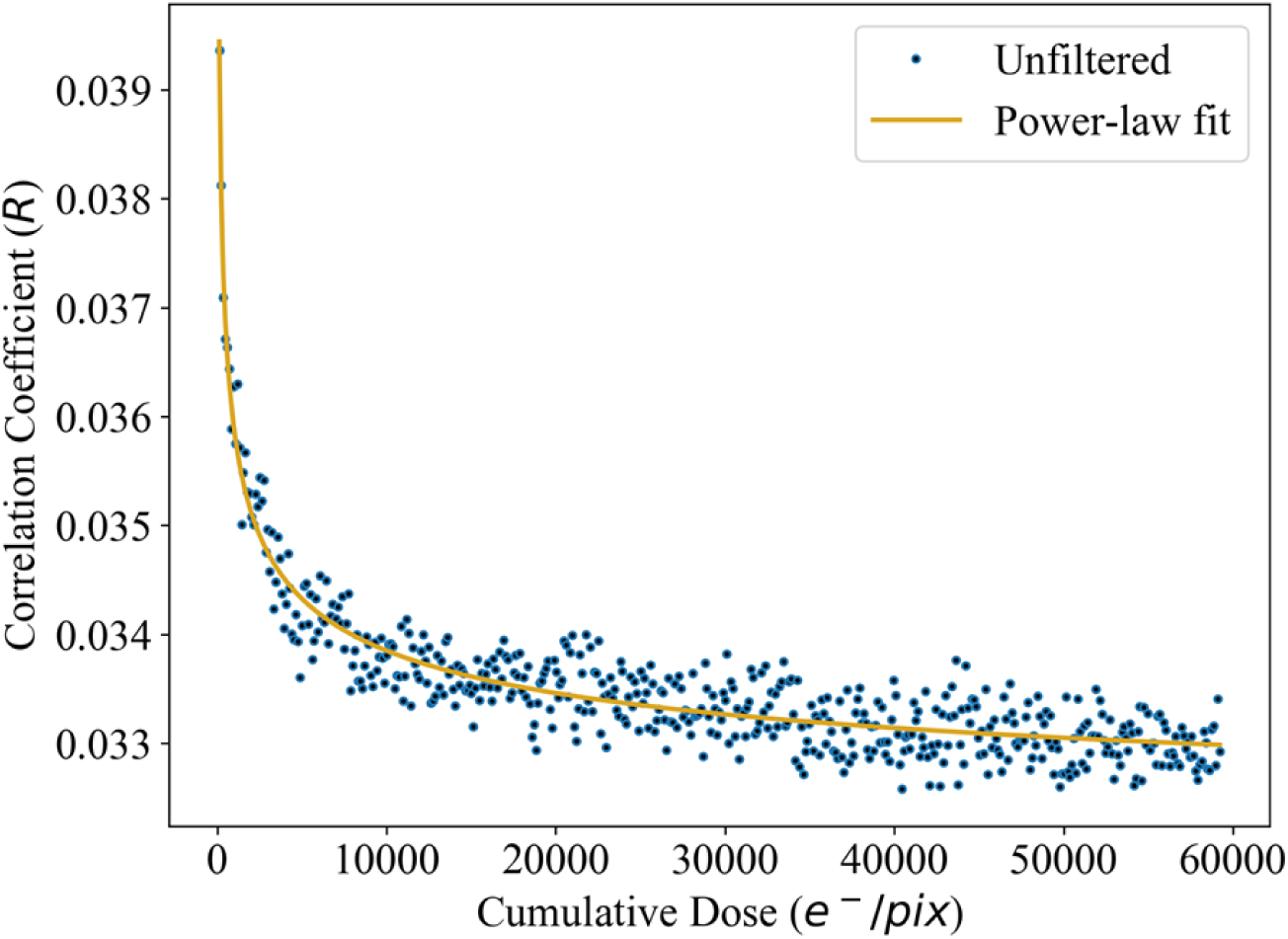
Pearson correlation for flat-field illuminated images shows a rapid-drop in adjacent frame correlation after a critical dose of approximately 4000 e-. A similar analysis of practical cryo-EM data shows no obvious drop off with dose as the image contrast dominates the signal.

Given the power-law decay in correlated noise with dose separation, from the perspective then of gain normalization, it is hypothesized that using the entire data set as in Afanasyev et al. is unnecessary.

Rather, a moving average filter with a radius ±*a* stacks may be sufficient, which would render the approach much more compatible with pipeline processing. In this approach all stacks have uniform weights, which greatly simplifies computational requirements. The moving average must skip, or ‘knock-out,’ the stack undergoing normalization from the moving-average, to avoid intentionally subtracting part of the image from itself. Then the calculation of each knock-out moving average (KOMA) filter requires two image adds (of the previous stack and the latest stack) and two subtractions (of the current stack and the last stack in the series), which is computationally inexpensive. End-points (at the start or end of data acquisition) are asymmetrically adjusted to maintain the same number of stack in the filter, so as to maintain an even signal-to-noise ratio. No gamma normalization has been applied in this case.

The effectiveness of the KOMA filter drops off near reciprocally towards an asymptotic limit. For both vacuum references and cryo-EM data, the correlated noise from KOMA filters with a radius of *a* are shown in Fig. 12(a), and the outlier filter combined with KOMA in Fig. 12(b). For KOMA alone with the cryo-EM data, the correlated noise is reduced below the baseline unfiltered stacks after subtraction of the mean of the 8 trailing and following frames, whereas for vacuum reference images only 4 stacks were required. The asymptotic limit for the reduction in correlated noise was 300 % for the cryo case and 650 % for the vacuum reference case. The power-law best fits to the data are,

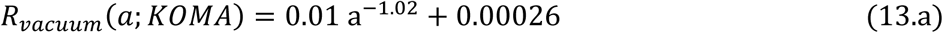

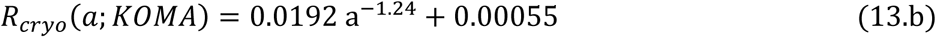

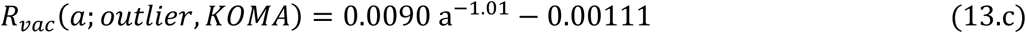

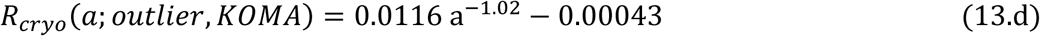

**Figure 12:**
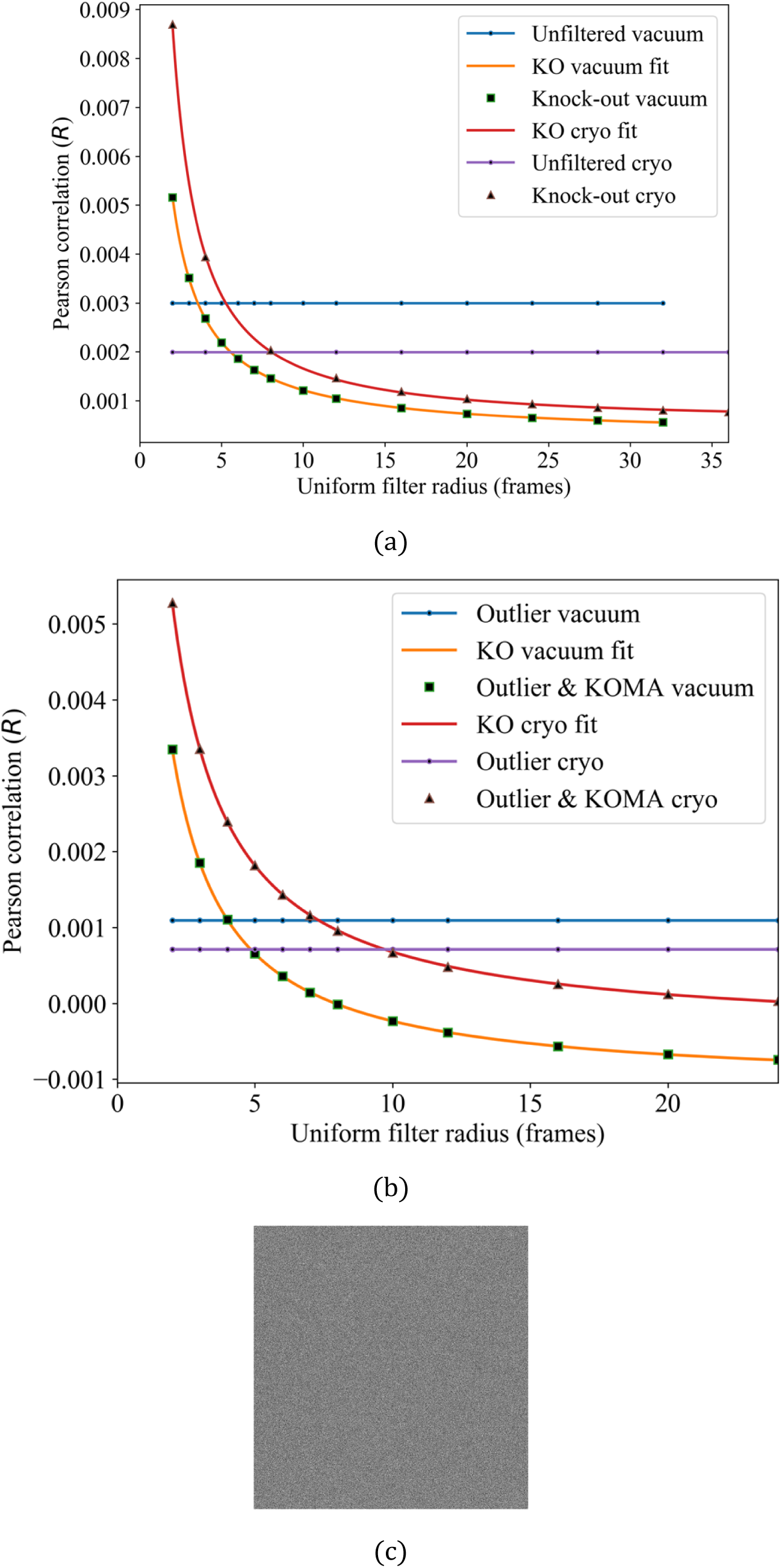
Analysis of correlated noise with the KOMA filter. (a) The correlated noise within frames of image stacks drops with a near-reciprocal relationship with the KOMA filter radius for both vacuum references and sample cryo-EM data. (b) When the outlier and KOMA filters are combined the correlated noise drops rapidly to zero before becoming, in the vacuum case, slightly anti-correlated with increasing KOMA radius. (c) After outlier and KOMA filtering with a 12-stack radius there is no visible detector artifacts remaining in a vacuum reference, even at a dose rate of *7e^-^/pix/s*.

which are effectively reciprocal functions in all but the cryo case with KOMA alone. When the outlier and KOMA filters are combined the correlated noise actually becomes anti-correlated as the radius of the KOMA filter increases, which likely indicates that the KOMA filter is too broad and the extra frames added no longer have significantly correlated artifacts with the stack under examination. In comparison the anti-correlation is not nearly as significant for the cryo-EM data.

The average dose rate for the cryo-EM data was 4.5*/pix/s* and *7e^−^/pix/s* for the vacuum references. Thus there is also an expectation of some coincidence counting errors in the vacuum reference case (Li et al., 2013).

Note that here the correlation coefficients are calculated amongst frames inside an individual stack after normalization, whereas in Fig. 11 the correlations were between stack sums. Therefore there is more random-nose in-between each correlation in Fig. 12(a) and as such the observed coefficients are lower. One must be careful not to over-interpret the cryo-EM results as the appearance of a minimum value in the Pearson coefficient may be indicative of a minimum in image contrast rather than correlated noise. In general, there appears to be little advantage to averaging more than ±25 stacks in the KOMA filter in the cases studied here. One must be cautious to avoid including high-contrast micrographs (such as those with carbon film or crystalline ice), however, and here the micrographs were pruned by hand. Automated detection of such high-contrast micrographs remains a subject of future work.

## 6 Conclusion

The introduction of fast, large-pixel count direct electron detectors has moved the field of electron microscopy inside the domain of “Big Data” in terms of data processing and storage requirements. Here an extension of the MRC file format is presented that permits on-the-fly data compression to lessen both transmission and storage requirements, using the meta-compression library *blosc*. *Blosc* is well suited as a library for Big Data purposes because it does not explicitly endorse any particular algorithm and intends to support new compression methods as they are developed. Development of a *blosc2* standard, with additional features, is currently underway. Also *blosc is* especially targeted towards high-speed compression codecs. With high-speed compression using the *zStandard* codec, file input-output rates are accelerated and archival storage requirements are reduced. The primary disadvantage of the use of a compressed MRC format is backward compatibility, which is mitigated by the availability of a command-line conversion tool. The simple implementation of the C-MRCZ library should facilitate its insertion into legacy codebases.

With the use of compression, sparse data can be compressed to very high ratios. Hence, data can be recorded in smaller dose fractions with a less-than-linear increase in data size. For very small dose fraction applications, such as cryo-electron tomography, electron crystallography, or software electron counting schemes, compression can reduce data transmission and storage requirements by 10–50x.

Effective compression requires the raw data to be stored as integers, which in turn requires artifact removal and gain normalization to be performed at the destination at least as well as vendors’ methods. The generation of outlier pixels in a direct-electron detector is a dynamic process and an algorithm which can effectively detect and suppress outliers was presented, reducing the correlated noise greater than five-fold over a simple gain normalization. The remaining degree of correlated noise likely sources from the CMOS electronics and active layer etching pattern. Here, an *a priori* approach of applying a gamma correction to the gain reference, and an *a posteriori* approach known as a knock-out moving average (KOMA) filter that is compatible with pipeline processing were presented. The gamma correction suppresses the etching but not the CMOS pattern, whereas the KOMA filter suppresses both. The outlier pixel and KOMA filters may be combined, in which case the degree of correlated noise can be negligible, even for relatively high dose-rates ~*7e^−^/pix/s*, where some coincidence counting loss is expected.

## Acknowledgements

The authors would like to thank the authors of the *blosc* library, Francesc Alted and Valentin Haenel, for their work on the library and feedback during our testing of MRCZ. The authors also thanks Nicolas Taylor for the cryo-TEM data used for characterization of the KOMA filter.

## Abbreviations

uint8: unsigned 8-bit integer computer data, range 0–255
float32: floating-point 32-bit computer data, ~6 significant figures
Blosc: blocked, shuffle, compress library
CDF: cumulative density function
CMOS: complementary metal-oxide semiconductor
PDF: probability density function
GB: Gigabyte
Gb: Gigabit
MB: Megabyte
KOMA: knock-out moving average

## References

Alted, F., 2014. Blosc, an extremely fast, multi-threaded, meta-compressor library [WWW Document]. Blosc Main Page. URL http://www.blosc.org/index.html (accessed 3.13.17).

Afanasyev, P., Ravelli, R.B.G., Matadeen, R., Carlo, S.D., Duinen, G. van, Alewijnse, B., Peters, P.J., Abrahams, J.-P., Portugal, R.V., Schatz, M., Heel, M. van, 2015. A posteriori correction of camera characteristics from large image data sets. Scientific Reports 5, 10317. doi:10.1038/srep10317

Cheng, A., Henderson, R., Mastronarde, D., Ludtke, S.J., Schoenmakers, R.H.M., Short, J., Marabini, R., Dallakyan, S., Agard, D., Winn, M., 2015. MRC2014: Extensions to the MRC format header for electron cryo-microscopy and tomography. Journal of Structural Biology, Recent Advances in Detector Technologies and Applications for Molecular TEM 192, 146–150. doi:10.1016/j.jsb.2015.04.002

Crowther, R.A., Henderson, R., Smith, J.M., 1996. MRC Image Processing Programs. Journal of Structural Biology 116, 9–16. doi:10.1006/jsbi.1996.0003

Duda, J., 2013. Asymmetric numeral systems: entropy coding combining speed of Huffman coding with compression rate of arithmetic coding. arXiv:1311.2540 [cs, math].

Duda, J., Tahboub, K., Gadgil, N.J., Delp, E.J., 2015. The use of asymmetric numeral systems as an accurate replacement for Huffman coding, in: 2015 Picture Coding Symposium (PCS). Presented at the 2015 Picture Coding Symposium (PCS), pp. 65–69. doi:10.1109/PCS.2015.7170048

Standard ECMA-404 [WWW Document], n.d. URL http://www.ecma-international.org/publications/standards/Ecma-404.htm (accessed 3.11.17).

Grant, T., Grigorieff, N., 2015. Measuring the optimal exposure for single particle cryo-EM using a 2.6 Å reconstruction of rotavirus VP6. eLife Sciences e06980. doi:10.7554/eLife.06980

Google Cloud Storage Pricing | Cloud Storage Documentation [WWW Document], n.d. Google Cloud Platform. URL https://cloud.google.com/storage/pricing (accessed 3.11.17).

Haenel, V., 2014. Bloscpack: a compressed lightweight serialization format for numerical data. arXiv:1404.6383 [cs].

HDF5 File Format Specification Version 3.0 [WWW Document], n.d. URL https://support.hdfgroup.org/HDF5/doc/H5.format.html (accessed 1.3.17).

Li, X., Mooney, P., Zheng, S., Booth, C.R., Braunfeld, M.B., Gubbens, S., Agard, D.A., Cheng, Y., 2013. Electron counting and beam-induced motion correction enable near-atomic-resolution single-particle cryo-EM. Nat Meth 10, 584–590. doi:10.1038/nmeth.2472

Li, X., Zheng, S.Q., Egami, K., Agard, D.A., Cheng, Y., 2013. Influence of electron dose rate on electron counting images recorded with the K2 camera. Journal of Structural Biology 184, 251–260. doi:10.1016/j.jsb.2013.08.005

lz4/lz4 [WWW Document], n.d. GitHub. URL https://github.com/lz4/lz4 (accessed 3.13.17).

Mastronarde, D.N., 2005. Automated electron microscope tomography using robust prediction of specimen movements. Journal of Structural Biology 152, 36–51. doi:10.1016/j.jsb.2005.07.007

McLeod, R.A., Kowal, J., Ringler, P., Stahlberg, H., n.d. Robust image alignment for cryogenic transmission electron microscopy. Journal of Structural Biology. doi:10.1016/j.jsb.2016.12.006

MessagePack: It's like JSON. but fast and small. [WWW Document], n.d. URL http://msgpack.org/index.html (accessed 3.11.17).

Welch, T.A., 1984. A Technique for High-Performance Data Compression. Computer 17, 8–19. doi:10.1109/MC.1984.1659158

Zheng, S.Q., Palovcak, E., Armache, J.-P., Verba, K.A., Cheng, Y., Agard, D.A., 2017. MotionCor2: anisotropic correction of beam-induced motion for improved cryo-electron microscopy. Nat Meth advance online publication. doi:10.1038/nmeth.4193

facebook/zstd [WWW Document], n.d. GitHub. URL https://github.com/facebook/zstd (accessed 3.13.17).

